# A single-dose MCMV-based vaccine elicits long-lasting immune protection in mice against distinct SARS-CoV-2 variants

**DOI:** 10.1101/2022.11.25.517953

**Authors:** Kristin Metzdorf, Henning Jacobsen, Yeonsu Kim, Luiz Gustavo Teixeira Alves, Upasana Kulkarni, Kathrin Eschke, M. Zeeshan Chaudhry, Markus Hoffmann, Federico Bertoglio, Maximilian Ruschig, Michael Hust, Maja Cokarić Brdovčak, Jelena Materljan, Marko Šustić, Astrid Krmpotić, Stipan Jonjić, Marek Widera, Sandra Ciesek, Stefan Pöhlmann, Markus Landthaler, Luka Čičin-Šain

**Affiliations:** Department of Viral Immunology, Helmholtz Centre for Infection Research, Braunschweig, Germany; Berlin Institute for Medical Systems Biology (BIMSB), Max Delbrück Center for Molecular Medicine in the Helmholtz Association (MDC), Berlin, Germany; Institute for Biology, Humboldt-Universität zu Berlin, Berlin, Germany; Infection Biology Unit, German Primate Center – Leibniz Institute for Primate Research, Göttingen, Germany; Faculty of Biology and Psychology, Georg-August-University Göttingen, Göttingen, Germany; Department of Medical Biotechnology, Institute for Biochemistry, Biotechnology and Bioinformatics, Technische Universität Braunschweig, Braunschweig, Germany; Center for Proteomics, University of Rijeka, Faculty of Medicine, Rijeka, Croatia; Department of Histology and Embryology, University of Rijeka, Faculty of Medicine, Rijeka, Croatia; Institute for Medical Virology, University Hospital Frankfurt, Goethe University Frankfurt, Frankfurt am Main, Germany; Fraunhofer Institute for Translational Medicine and Pharmacology ITMP, Frankfurt am Main, Germany; German Centre for Infection Research (DZIF), External partner site Frankfurt, Germany; Centre for Individualized Infection Medicine (CiiM), a joint venture of HZI and MHH, Hannover, Germany

**Keywords:** SARS-CoV-2, COVID-19, MCMV, vaccination, single-dose, long-lasting protection, mouse, *in vivo*

## Abstract

Current vaccines against COVID-19 elicit immune responses that are overall strong but wane rapidly. As a consequence, the necessary booster shots have led to vaccine fatigue. Hence, vaccines that would provide lasting protection against COVID-19 are needed, but are still unavailable. Cytomegaloviruses (CMV) elicit lasting and uniquely strong immune responses. Used as vaccine vectors, they may be attractive tools that obviate the need for boosters. Therefore, we tested the murine CMV (MCMV) as a vaccine vector against COVID-19 in relevant preclinical models of immunization and challenge. We have previously developed a recombinant murine CMV (MCMV) vaccine vector expressing the spike protein of the ancestral SARS-CoV-2 (MCMV^S^). In this study, we show that the MCMV^S^ elicits a robust and lasting protection in young and aged mice. Notably, S-specific humoral and cellular immunity was not only maintained but even increased over a period of at least 6 months. During that time, antibody avidity continuously increased and expanded in breadth, resulting in neutralization of genetically distant variants, like Omicron BA.1. A single dose of MCMV^S^ conferred rapid virus clearance upon challenge. Moreover, MCMV^S^ vaccination controlled two immune-evading variants of concern (VoCs), the Beta (B.1.135) and the Omicron (BA.1) variants. Thus, CMV vectors provide unique advantages over other vaccine technologies, eliciting broadly reactive and long-lasting immune responses against COVID-19.

**Authors Summary:** While widespread vaccination has substantially reduced risks of severe COVID presentations and morbidity, immune waning and continuous immune escape of novel SARS-CoV-2 variants have resulted in a need for numerous vaccine boosters and a continuous adaptation of vaccines to new SARS-CoV-2 variants. We show in proof of principle experiments with a recombinant murine cytomegalovirus expressing the SARS-CoV-2 spike protein (MCMVS) that one immunization with a CMV vaccine vector drives enduring protection in both young and aged mice, with long-term maturation of immune responses that broaden the antiviral effects over time. Hence, this approach resolves issues of immune waning and mitigates the effects of COVID-19 evolution and immune escape, reducing the need for additional immunizations and potentially improving vaccine compliance.

## Introduction

The corona-virus disease 2019 (COVID-19), caused by the severe acute respiratory syndrome coronavirus 2 (SARS-CoV-2) (1), had massive impact on public health. The virus rapidly spread across multiple countries in 2020, leading to millions of confirmed cases and deaths in the subsequent years (2). The elderly have been particularly affected by SARS-CoV-2, with a higher risk of severe COVID-19. The exhibited little genetic variation during the early months of the pandemic, with only a single spike substitution (D614G) in the PANGO lineage B.1, which rapidly became dominant in Europe in late 2020 (3). New variants emerged in 2021, characterized by an increasing number of mutations in the spike (S) protein sequence and immune escape potential, including the Beta (B.1.351) (4), the Delta (B1.617.2) (5), or the Omicron variant (B.1.1.529) (6) and its derivatives. Numerous COVID-19 vaccines that induce immune responses against the S glycoprotein have been developed and authorized for use with unprecedented speed. However, vaccine-induced immunity wanes within months, resulting overall in reduced protection against infections and disease (4, 7-9). In addition, emerging variants have been increasingly capable of escaping vaccine-induced immunity driving the need for continued vaccine adaptation and booster shots (10-12). Thus, acute (13) or long-term (14-16) effects after SARS-CoV-2 infection remain a public health issue and a vaccine that confers lasting and broad immune protection remains unmet.

Recently, our research group has demonstrated the capacity of a mouse cytomegalovirus (MCMV)-based COVID-19 vaccine candidate, encoding the S protein derived from the Wuhan prototype (MCMV^S^), elicits robust and persistent humoral as well as cellular immune responses in mice (17). MCMV is a well-characterized model virus for human CMV (HCMV) infection (18). CMVs are strictly species-specific for their cognate host, and induce robust anti-viral immunity, which persists over a lifetime (19-22). This long-term maintenance of antigen-specific lymphocytes, known as memory inflation (22, 23), is characterized by an accumulation and persistence of CD8 T cells with an effector memory (TEM) phenotype, recognizing immunodominant viral antigens (23, 24). Therefore, transgenic antigens from other infectious agents have been expressed in recombinant MCMVs (25), and such vaccine vectors were shown to provide long-term immune protection (26). This concept was validated in other models of CMV immunization, where a peculiarly strong CD8 T cell immunity was crucial for immune protection (26-31). More recently it was noticed that MCMV infection also results in lasting humoral immune responses toward viral antigens (32), which was exploited to retarget immune protection against other viruses (33), including influenza (17).

The S protein of the original strains of SARS-CoV-2 does not bind to mouse ACE2 (mACE2) receptors. Hence, mouse models of SARS-CoV-2 infection were initially restricted to transgenic mice expressing the human ACE2 receptor (hACE2), or mouse-adapted SARS-CoV-2 variants. However, the naturally occurring N501Y mutation in the S protein is sufficient for mACE2 binding (34). Since this mutation was present in various variants of concern (VoCs), including Beta and Omicron, these variants could also be used in infection experiments with non-transgenic mice (35).

Here, we show that immunization with MCMV^S^ elicited virus neutralizing titers (VNTs) that increased for at least 6 months, concomitant with an increase of antibody avidity and resulting in a broadening of neutralization effects to genetically distant variants. The immunization protected vaccinated K18-hACE2 mice against lethality and disease manifestations upon challenge with D614G, or Delta VoC. Furthermore, the same vaccination protected BALB/c mice against challenge with the immune-evasive Beta and Omicron strains, and the protection lasted for at least 5 months. Hence, our results argue that CMV-based SARS-CoV-2 vaccines may provide long-term and broad protection against multiple VoCs.

## Results

### Protection against SARS-CoV-2 D614G elicited by a single dose of MCMV^S^ vaccination

We have previously shown that a single dose of a MCMV-based vaccine vector encoding the S-antigen (MCMV^S^) from SARS-CoV-2 triggers a robust inflationary CD8 T cell response against the S-encoded octameric peptide VNFNFNGL in C57BL/6 mice and neutralizing antibody responses which persisted in BALB/c mice for a minimum of 90 days (17). Now we immunized K18-hACE2 mice, which express the human ACE2 receptor (36) and thus may be infected by SARS-CoV-2. We used MCMV^S^, a vector control (MCMV^WT^), or PBS for immunization/mock treatment and challenged the mice with a lethal dose of SARS-CoV-2 B.1 (D614G), 6 or 12 weeks later (Fig. 1A).

**Figure 1:**
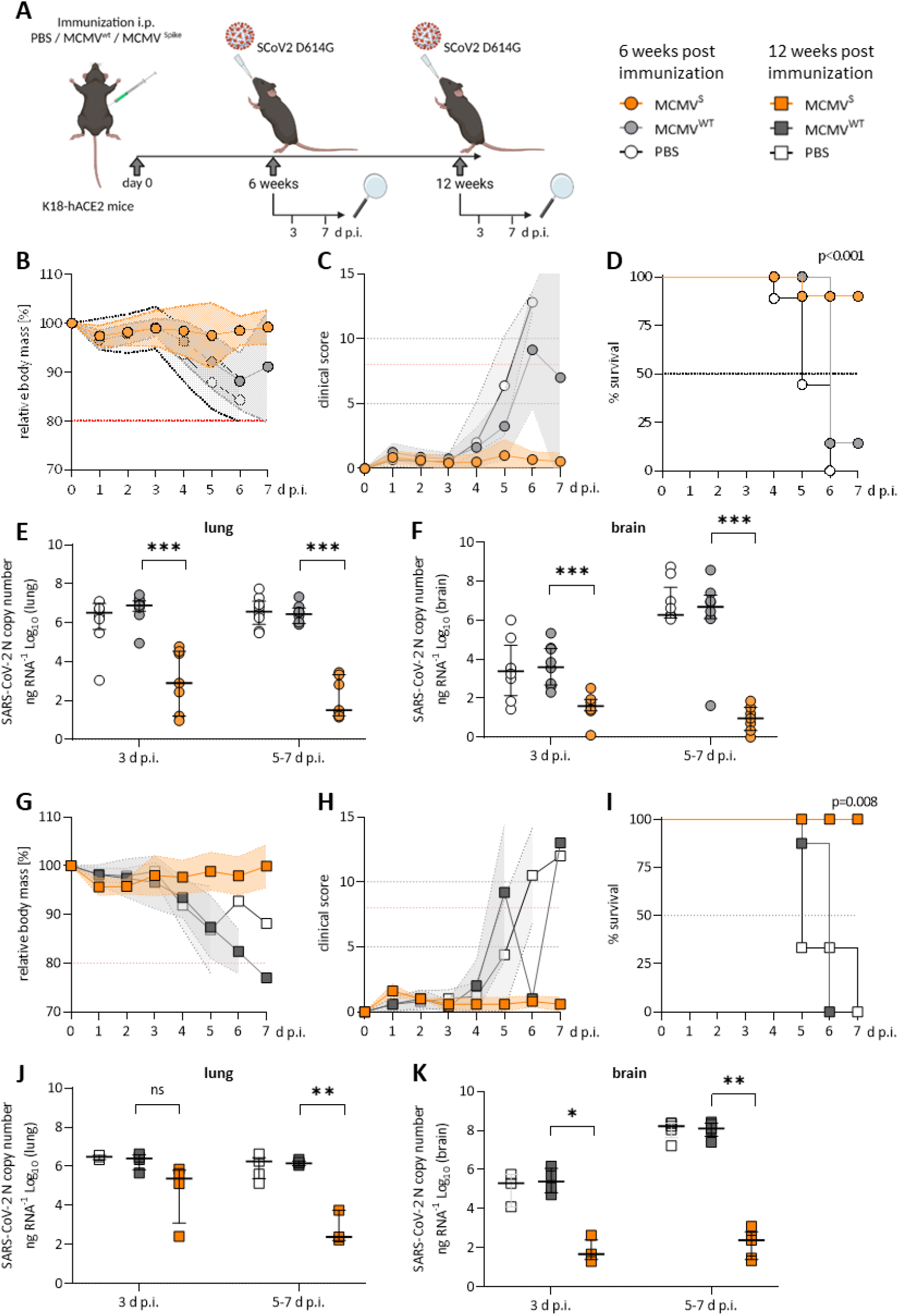
Protection against SARS-CoV-2 D614G elicited by a single dose of MCMV^S^ vaccination. **(A)** Schematic image of the experimental setup and legend for panels B-K. Created with BioRender.com **(B-F)** Mice were challenged with 2×10^3^ PFU of SARS-CoV-2 D614G 6 weeks or **(G-L)** 12 weeks post immunization. **(B, G)** Percentage of body mass relative to day 0 and **(C, H)** daily clinical scores based on appearance, activity and posture upon challenge are shown. The red dotted line indicates the thresholds resulting in a humane end-point which were used as proxies for survival. **(D, I)** Mock-survival kinetics of challenged mice representing the Percentage [%] of mice reaching a humane end point. A log-rank (Mantel-cox) test was used for the statistical analysis. **(E-F, K-L)** SARS-CoV-2 N gene copy numbers as copy number/ng RNA Log10 at 3 d p.i. and 5-7 d p.i. with SARS-CoV-2 D614G in lung homogenates **(E, K)** and brain homogenates **(F, L)**. Pooled data (n=6-8 per group) from two independent experiments are shown. Statistical significance for was calculated using Welch’s t test (two-tailed) (* p < 0.05, ** <u>p</u> <0.01, *** p < 0.001).

To quantify the CD8+ T cell response, blood was harvested before the SARS-CoV-2 challenge at 1 or 3 months post-immunization, and blood leukocytes were analyzed by flow cytometry (Supplementary Fig. 1). MCMV^S^ and MCMV^WT^ elicited comparable frequencies of CD8 T cells in total, primed, effector (T_EFF_), terminally differentiated effector (T_TDE_), effector memory (T_EM_), and central memory (T_CM_) compartments (Supplementary Fig. 1A). However, only MCMV^S^-vaccinated mice displayed high frequencies of VNFNFNGL-specific CD8+ T cells (Supplementary Fig. 1B). The frequency of S-specific T cells was approximately two-fold increased at 3 months over 1 month post-immunization in all tested subsets (Supplementary Fig. 1B) and this was confirmed by dynamic monitoring of S-specific CD8 T cells in the blood (Supplementary Fig. 1C). Likewise, neutralizing titers were typically higher at 3 than at 1 month post-immunization (Supplementary Fig. 1D), in line with our previous report in non-transgenic mice (17). We challenged these mice with 2×10^3^ PFU of SARS-CoV-2 at the indicated time points (Fig. 1A) and monitored weight loss and clinical scores to evaluate protection against COVID-19. Immunized animals that robustly responded to vaccination (Supplementary Fig. 1) were protected against challenge at 6 weeks, as evidenced by maintained weight (Fig. 1B) and by the absence of clinical signs (Fig. 1C), while most mock-vaccinated mice became severely ill, reaching the humane end-point by day 5 post-infection (d p.i.) or earlier (Fig. 1D). At 3 and at 5-7 d p.i., MCMV^S^ vaccination decreased the median SARS-CoV-2 viral RNA load in lungs and brains by a factor of 1000 or more relative to mock-vaccinated groups (Fig. 1E-F). The results were similar in the trachea, stomach, heart and spleen (Supplementary Fig. 2A). The immunity of clinically approved vaccines against COVID-19 gradually wanes over time (37, 38). To assess if the protective effects of MCMV^s^ is reduced over time, mice were challenged at 3 months post immunization. All MCMV^S^-vaccinated mice were protected against SARS-CoV-2 infection, in terms of body mass reduction (Fig. 1G), clinical signs (Fig. 1H) and survival (Fig. 1I). Viral RNA burdens were reduced in the MCMV^S^-immunized group in all examined organs (Fig. 1 J-K and Supplementary Fig. 2B). Some MCMV^S^-vaccinated mice presented viral RNA in lungs at 3 d p.i., but only minor amounts of viral RNAs were detected in this organ at 7 d p.i. (Fig 1J), arguing that a single administration of the MCMV^S^ vaccine inhibits virus replication. Moreover, the viral RNA load in the brain was ∼10,000-fold higher in the controls compared to the MCMV^S^ immunized mice at day 5-7 post-challenge (Fig. 1K). In sum, single-dose MCMV^S^ immunization protected animals from COVID-19 at 6 and 12 weeks post-vaccination.

### Sustained neutralizing antibody responses against SARS-CoV-2 Beta and Omicron variants upon MCMV^S^ immunization

We next tested MCMV^s^ elicited immunity against the immune evasion Beta (B.1.351) and Omicron (BA.1) VoC. Since both of these VoCs contain the N501Y mutation in the S protein, SARS-CoV-2 infection is possible in non-transgenic mice, such as BALB/c and C57BL/6 (34). The spike used in MCMV^S^ corresponds to the Wuhan variant of SARS-CoV-2, and therefore does not engage the mACE2 receptor in such mice, potentially resulting in fewer antigen-specific side effects. To assess whether MCMV^S^ can protect BALB/c in the long term, we challenged mice at five months post-vaccination with the Beta or the Omicron BA.1 variant. A short-term cohort was challenged at 6 weeks after immunization as a point of reference (Fig. 2A). In the short-term cohort, MCMV^S^-immunized animals retained their body mass and displayed minimal clinical symptoms, while MCMV^WT^ mock-immunized animals lost approximately 10% of their initial body mass at day three post-infection (Fig. 2B-C). Irrespective of the immunization status, animals that were challenged with the Omicron BA.1 variant showed no reduction in body mass and overall very mild clinical symptoms (Supplementary Fig. 3A-B), in line with reports by other studies.

**Figure 2:**
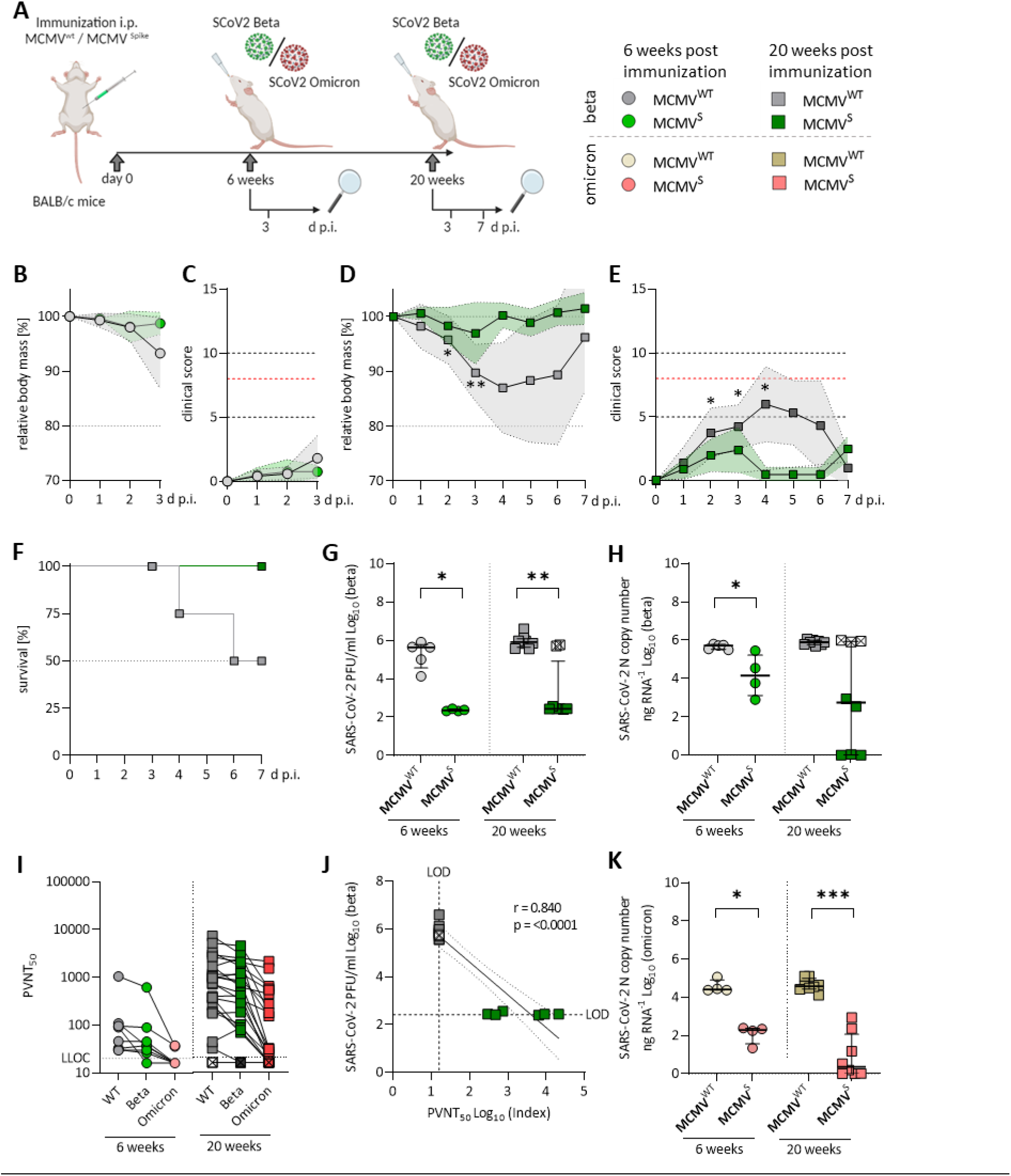
Sustained neutralizing antibody responses against Beta and Omicron BA.1 variants upon MCMV^S^ immunization. **(A)** Schematic image of the experimental setup and legend. Mice were challenged with 6×10^4^ PFU of SARS-CoV-2 in all settings. Created with BioRender.com. **(B, C)** Mice were challenged at 6 weeks after immunization. Relative body mass **(B)** and clinical scores **(C)** were monitored daily upon challenge with the Beta variant. **(D-F)** Mice were challenged with SARS-CoV-2 Beta 20 weeks after immunization. **(D)** Percentage of body mass, **(E)** daily clinical scores and **(F)** survival kinetics of challenged mice represented as the percentage of mice reaching the humane end point due to a high clinical score. The threshold resulting in this humane end-point is indicated by the red dotted line in panel **(E). (G)** Infectious virus titers in lungs of SARS-CoV-2 Beta-challenged mice 6 weeks (left) and 20 weeks (right) post immunization. **(H)** Relative viral RNA loads in lungs of SARS-CoV-2 Beta-challenged mice normalized to GAPDH. **(J)** Pseudovirus serum neutralization titers resulting in a 50% reduction of SARS-CoV-2 infection (PVNT_50_). PVNT_50_ at 6 weeks (n=8; left) and 20 weeks (n=24; right) post-vaccination against the Index (Wuhan), Beta and Omicron BA.1 variants. Titers were assessed seven days before SARS-CoV-2 challenge. Dot plots representing the same mice are connected by lines. Solid horizontal lines show the median and the dotted line indicates the detection limit. **(J)** Correlation of Infectious virus titers in lungs and PVNT50 against the Index strain of SARS-CoV-2 Beta-challenged mice after 20 weeks post immunization. **(K)** Relative viral RNA loads in lungs of SARS-CoV-2 Omicron BA.1-challenged mice normalized to GAPDH. Statistical significance for was calculated using Welch’s t test (two-tailed) (* p < 0.05, ** p <0.01, *** p < 0.001).

At five months post-vaccination, MCMV^S^-vaccinated animals remained protected against clinical disease upon SARS-CoV-2 Beta challenge (Fig. 2D and E). A randomly selected subset of animals from the long-term cohort was additionally monitored for seven d p.i. for clinical signs and survival. Half of the animals in the mock-immunized group reached the humane end-point, while this occurred in none of the immunized animals (Fig. 2F). Challenge with Omicron BA.1 resulted in no discernible body mass reduction in any of the groups and mild disease in mock-immunized animals 6 weeks (Supplementary Fig. 3A) and 20 weeks post MCMV-immunization (Supplementary Fig. 3B). Altogether, our data indicate that a single dose of MCMV^S^ vaccination provided a robust and lasting protection against the antigenically distinct Beta variant of SARS-CoV-2, while clinical protection against Omicron BA.1 was hard to assess due to overall low pathogenicity of this variant in mice.

Vaccine efficacy was further assessed by measuring viral RNA copies and infectious titer in the lungs at 3 d p.i.. Both short- and long-term MCMV^S^-immunized animals harbored significantly fewer infectious particles in the lungs than the control groups (Fig. 2G). Interestingly, viral RNA load in the short-term immunized cohort was on average slightly higher than in the long-term cohort (Fig. 2H), which might indicate that antiviral activity improved with a longer immunization period. However, three animals in the 20-weeks post-vaccination group showed high viral RNA loads upon infection with the Beta VoC and two of the samples were also highly infectious in terms of PFUs (Fig 2G). In clinical settings, protection against infection and disease of COVID-19 is associated with the presence of neutralizing antibodies (nAb) targeting SARS-CoV-2 (39, 40). Hence, we tested the neutralizing capacity of antibodies targeting different variants of SARS-CoV-2 before and after Beta or Omicron BA.1 challenge. The nAb titers against the Index (Wuhan), Beta and Omicron BA.1 strains were higher at 20-weeks than at 6-weeks post-immunization (Fig. 2I). Importantly, the two immunized mice that showed high infectious virus titers were also the only two mice with undetectable nAb titers (Fig. 2J), indicating that immunization of these animals failed.

Importantly, MCMV^S^-vaccinated animals showed a strong capacity to control replication of the antigenically distinct Omicron BA.1 variant. We observed 100-fold reduced viral RNA loads in the short-term cohort and 10,000-fold reduced virus loads in the long-term challenge scenario (Fig. 2K). Infectious Omicron BA.1 virus remained undetectable in any scenario (data not shown).

Collectively, a receptor-inert S antigen delivered through an MCMV-based vector prompted robust and persistent nAb reactions, with no signs of immune waning and predictively correlated with a decrease in SARS-CoV-2 viral titers upon challenge.

### Single-dose of MCMV^S^ provides inflationary immunogenicity in aged mice

The elderly population is of special concern when it comes to COVID-19, due to their increased vulnerability and diminished vaccine efficacy. Hence, we assessed the potential of MCMV^S^-based vaccines in aged hosts. Specifically, we immunized aged K18-hACE2 mice at 14-16 months of ageeither with MCMV^S^ or the MCMV^WT^ as control vector (Fig. 3A). The neutralization capacity of sera was examined before SARS-CoV-2 challenge at 6 and 12 weeks, as well as up to 24 weeks postimmunization. Virus-nAb titers increased over time, albeit this increase was not significant for the Index or the Beta variant (Fig. 3B). Only a few mice showed detectable titres of neutralising antibodies against Omicron BA.1 or BA.4/5 shortly after immunization. However, cross-reactive titers against these variants increased significantly over time against both Omicron, with approximately half of the animals exhibiting robust titers 6 months post-vaccination (Fig. 3B).

**Figure 3:**
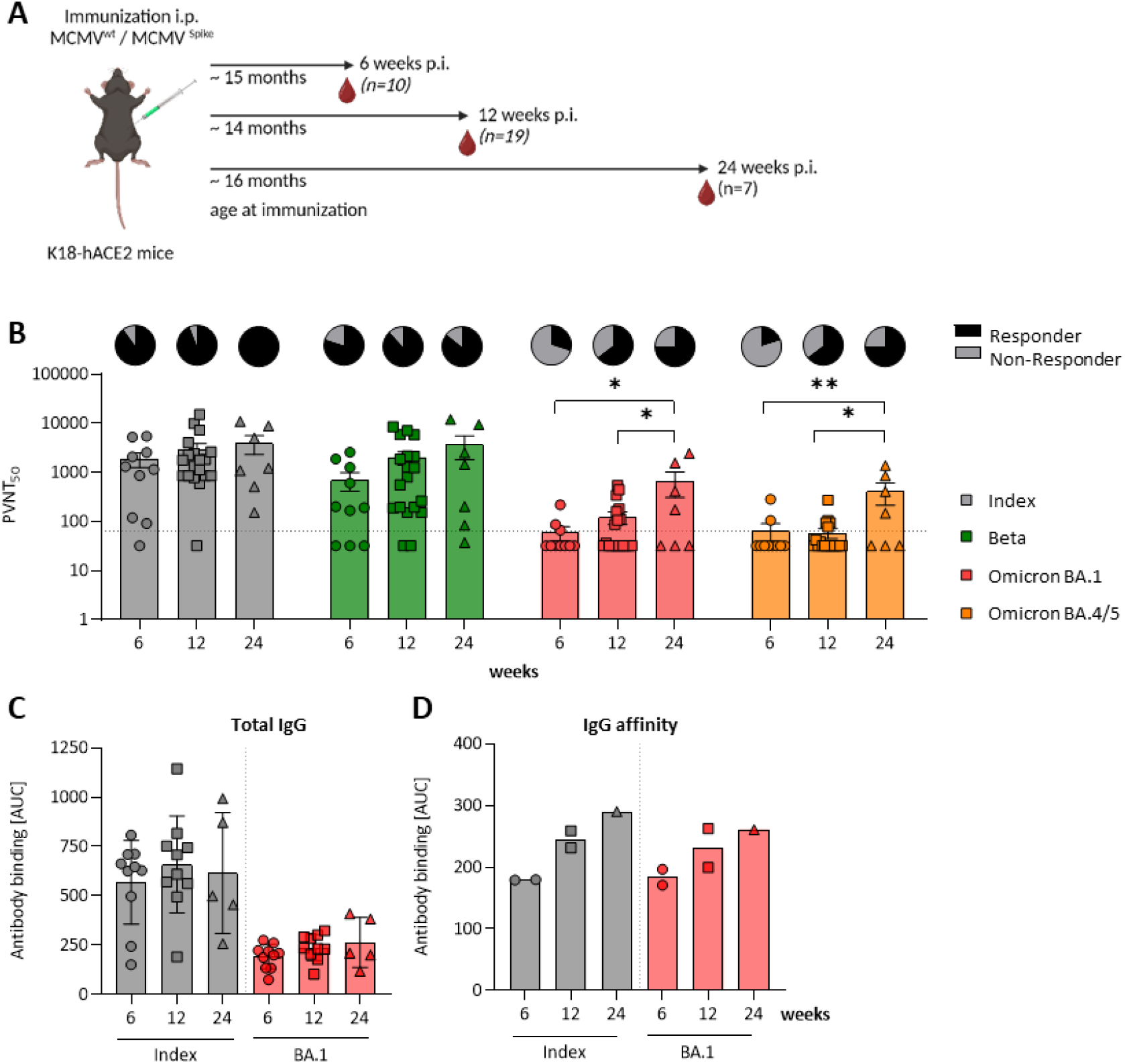
A single-dose of MCMV^S^ provides inflationary immunogenicity in aged mice. **(A)** Schematic image of the experimental setup. Created with BioRender.com. **(B)** Pseudovirus neutralization titers that resulted in a 50% of reduction of SARS-CoV-2 infection (PVNT_50_). Sera of 6 (n=10), 12 (n=19) and 24 (n=7) weeks MCMV^S^-immunized mice were tested against the Index (Wuhan; grey), Beta (blue), Omicron BA.1 (red) and Omicron BA.4/5 (orange) SARS-CoV-2 strains. Each symbol indicates an individual mouse. Solid horizontal lines show the median and the dotted line indicates the detection limit. Pie charts indicate the percentage [%] of animals, that responded (black) or non-responded (grey) with nAb titers. **(C-D)** SARS-CoV-2 Spike specific IgG response against the Index and Omicron BA.1 variants in murine sera after 6, 12, and 24 weeks post-immunization. **(C)** Total IgG and **(D)** IgG affinity measured by antibody binding [AUC] in presence of graded amounts of NaSCN using ELISA. Statistical significance for was calculated using Welch’s t test (two-tailed) (* p < 0.05, ** p <0.01).

After repeated antigen exposure, B cells undergo somatic hypermutation and affinity maturation processes, yielding high-affinity antibodies that efficiently bind to antigens, in our case SARS-CoV-2 S (41-43). Thus, re-exposure of primed B cells to the antigen improves virus neutralization efficacy due to affinity maturation. Since MCMV-based vaccines stimulated robust antibody responses with increasing neutralization capacity (17), we considered that these functional responses may be driven by affinity maturation, or by polyclonal B-cell expansion and a pure increase in antibody titers. Hence, we quantified SARS-CoV-2 S-specific total IgG levels upon vaccination over time (Fig 3C). Kinetic measurements revealed no significant differences in total IgG levels, against both the Index and the Omicron BA.1 variant (Fig. 3C). Serum samples from control mice that underwent mock immunization were typically below the detection threshold (data not shown). We next tested IgG binding avidity by exposing sera to various concentrations of sodium thiocyanate (NaSCN), as described previously (17). We observed an increase in avidity over time, from 6 and 12 to 24 weeks post-immunization (Fig. 3D). Consistent levels of total anti-S IgG alongside a concurrent rise in binding affinity within the same sera may indicate that the increase in neutralization was driven primarily by affinity maturation.

To identify the limits of our vaccine approach, we subjected it to a stress-test, where we analysed long-term immunity upon a vaccine dose that was 5-fold lower than in previous experiments. At 16 months post-immunization, spike-specific splenic CD8 T cells were detectable at sustained levels, with a median frequency of 10% of VNF-specific cells in the CD8 population (Supplementary Fig. 4A), and a predominant effector phenotype (Supplementary Fig. 4B). Furthermore, some mice still harboured nAb against SARS-CoV-2, including the BA.4/5 variant (Supplementary Fig. 4C), while anti-spike IgG could be observed by ELISA (Supplementary Fig. 4D). Thus, a single dose of the MCMV^S^-based vaccine provided lasting, potentially lifelong, humoral and cellular immunity in aged mice.

### Aged MCMV^S^ immunized mice show full and long-lasting protection against heterologous SARS-CoV-2 challenge

We subsequently assessed the immune protection provided to aged mice by a single-dose MCMV^S^ vaccination. We challenged aged K18-hACE2 mice with two SARS-CoV-2 variants. Mice were infected with 2×10^3^ PFU of SARS-CoV-2 Delta or Omicron BA.1 at 6 or 12 weeks post-immunization (Fig. 4A). Mice immunized with MCMV^S^ were fully protected against body mass loss (Fig. 4B) and clinical manifestations of disease (Fig. 4C) for up to 12 weeks post-immunization when challenged with the virulent Delta variant, while mock-vaccinated mice exhibited severe illness and approximately half among them reached the humane end-point by day five post-infection (Fig. 4D). Mice challenged with the Omicron BA.1 variant demonstrated overall minimal body mass loss (Fig. 4E) and exhibited only mild clinical symptoms (Fig. 4F), irrespective of immunization status, consistent with our previous findings (Fig. 2). Moreover, protection against body mass reduction (Supplementary Fig. 5A) and clinical signs (Supplementary Fig. 5B) was observed in animals challenged 6 weeks after MCMV^S^ immunization when compared to MCMV^WT^-controls and they consistently survived (Supplementary Fig. 5C). Vaccine efficacy was further evaluated by quantifying the viral RNA burden (Fig. 4H) and infectious titers in the lungs (Fig. 4I) and brain (Fig. 4J) at five d p.i.. Analyses of viral load in SARS-CoV-2 Delta-challenged mice revealed a >1000-fold reduction in lungs (Fig. 4H, *top*) and brains (Supplementary Fig. 5D) in immunized animals. We also observed a marked (∼100-fold) reduction of SARS-CoV-2 viral RNA in lungs (Fig. 4H, *bottom*) of mice challenged with the Omicron variant, while brains showed low levels of Omicron mRNA, even in mock-vaccinated mice (Supplementary Fig. 5E).

**Figure 4:**
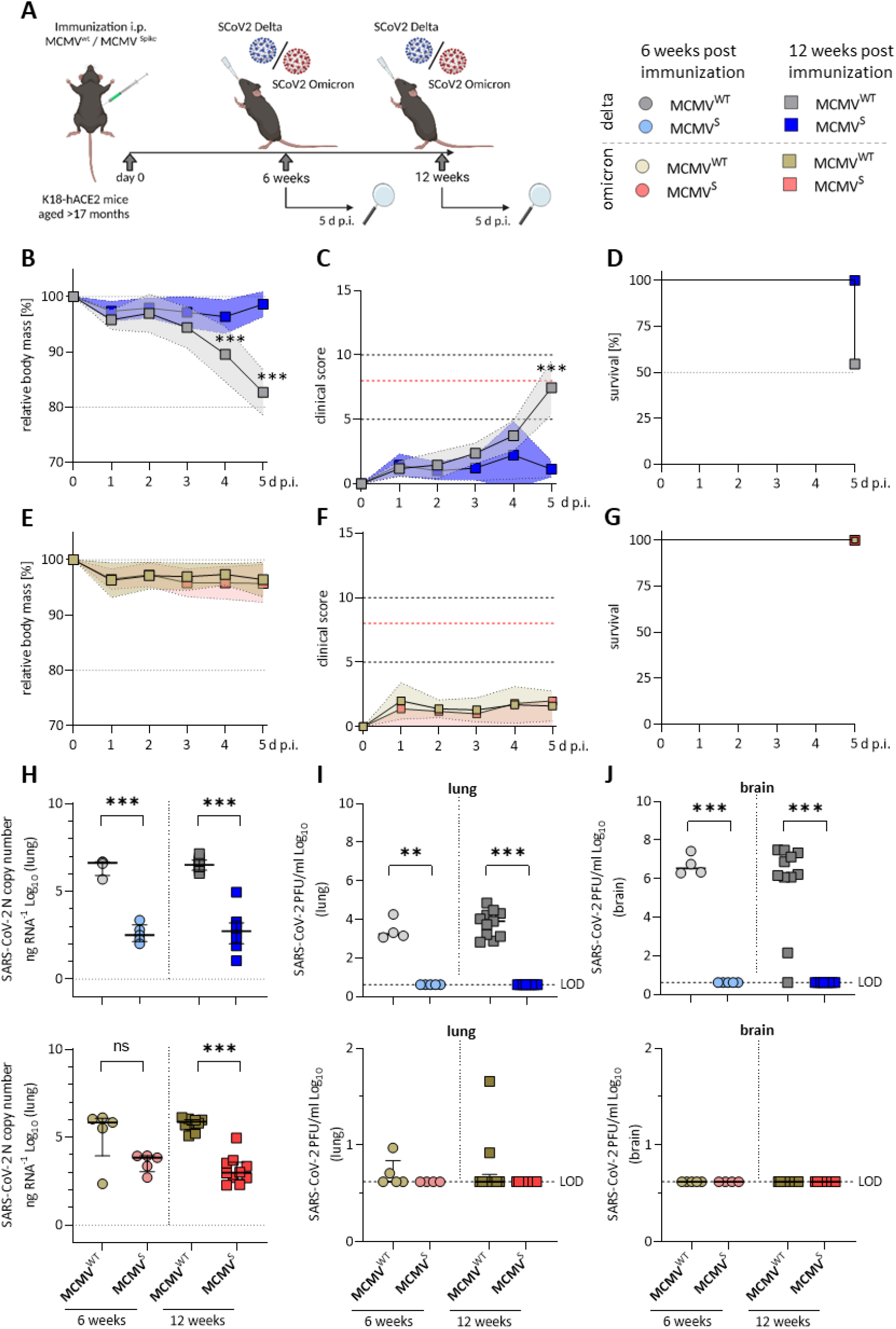
Aged MCMV^S^ immunized mice show full and long-lasting protection against heterologous SARS-CoV-2 challenge. **(A)** Legend of experimental cohorts. Created with BioRender.com. **(B-G)** Mice were challenged with 2×10^3^ PFU of SARS-CoV-2 **(B-D)** Delta or **(E-G)** Omicron BA.1 12 weeks after immunization. **(B, E)** Percentage of initial body mass and **(C, F)** daily clinical scores upon challenge are shown. The red dotted line indicates the threshold in score resulting in a humane end-point. **(D, G)** Survival kinetics of challenged mice representing the Percentage [%] of mice reaching a humane end point. A log-rank (Mantel-cox) test was used for the statistical analysis. **(H)** SARS-CoV-2 N gene copy numbers as copy number/ng RNA Log10 at 5 d p.i. with SARS-CoV-2 Delta (top) or Omicron BA.1 (bottom) in lung homogenates **(I-J)** Infectious virus titers in **(I)** lungs and **(J)** brains of SARS-CoV-2 Delta-(top) or Omicron BA.1 (bottom) challenged mice. Statistical significance was calculated using Welch’s <u>t</u> test (two-tailed) (* p < 0.05, ** p <0.01, *** p < 0.001).

MCMV^S^ immunized mice harbored no detectable infectious virus particles in lungs (Fig. 5I) and brains (Fig. 5J) at five days post challenge, indicating efficient and fast clearance in immunized animals. Taken together, MCMV^S^ fully protected aged mice against SARS-CoV-2 Delta challenge up to three months post-vaccination and reduced viral loads of the Delta and the antigenically distinct Omicron BA.1 variant.

**Figure 5:**
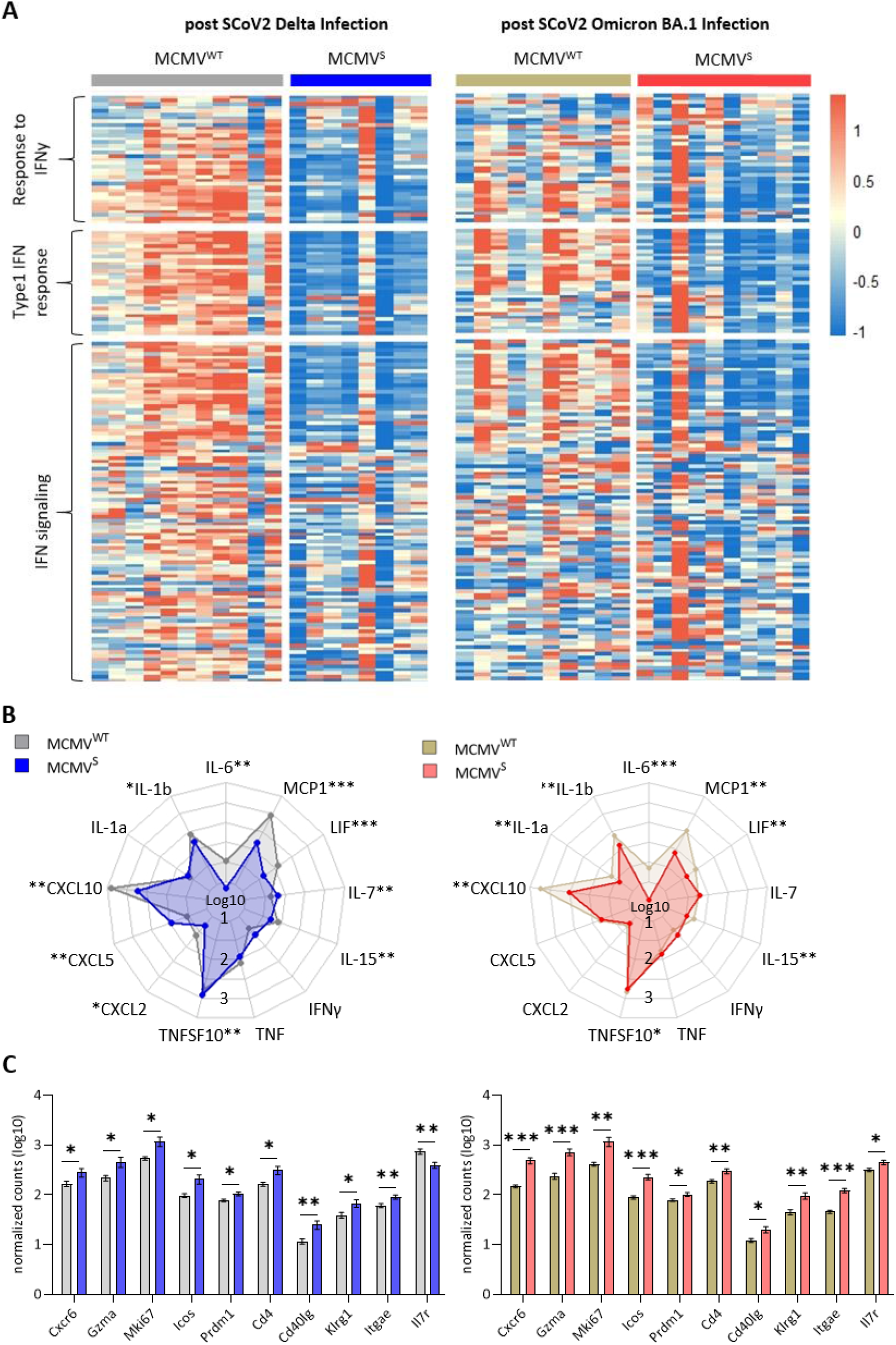
MCMV^S^ shows efficacy against the subclinical inflammation progression of Omicron BA.1. **(A)** Heat maps of differently expressed genes in response to vaccination involved in the response to IFNy, type 1 IFN response and related to IFN signaling (44). Differentially expressed genes of mice infected with the Delta (left) or Omicron variant (right) of SARS-CoV-2 and vaccinated with the MCMV^WT^ or S-modified MCMV vaccine was performed after bulk RNA sequencing of the lung tissue. Columns represent different animal samples and rows, genes. Shown are z-scores of DESeq2-normalized data and color scale ranges from red (10 % upper quantile) to blue (10 % lower quantile), showing up- or downregulation in expression of the selected genes, respectively. **(B)** Spider plot of differentially expressed cytokines between the 2 vaccine strategies. Values are DESeq2-normalized data in a log10 scale represented for each gene analyzed. **(C)** Heat map of differentially expressed genes involved in T cell activation and proliferation (45). Shown are z-scores of DESeq2-normalized data and color scale ranges from red (10 % upper quantile) to blue (10 % lower quantile), showing up- or downregulation in expression of the selected genes, respectively. N = 8-11 animals per group. (D) Cell-type deconvolution of bulk RNA-Sequencing performed with GEDIT for cell type prediction based on blood reference matrix provided by the program. Shown are fractions of the cell type selected of the total count.

### MCMV^S^ protects against subclinical inflammation following Omicron BA.1 infection

We used bulk RNA sequencing to investigate gene expression patterns associated with inflammation in lung samples after challenge with both Delta and Omicron variants in our long-term vaccination cohort. We observed a consistent difference in expression levels of genes involved in pathogen response and inflammation between the vaccinated mice and their mock vaccinated controls. The expression of genes related to interferon-gamma (IFNγ) and the type I interferon response, as well as Interferon (IFN) signalling cascade were clearly reduced in vaccinated mice upon both Delta and Omicron challenge (Fig. 5A). These data were in line with the reduction in symptoms observed upon Delta challenge and indicated that Omicron BA.1 infection induces a sub-clinical inflammation in mice, which was clearly reduced following vaccination. Cytokines that play crucial roles in inflammatory processes and are suspected to contribute to immunopathology were analyzed in more depth. Several among them including IL-6, CXCL10, IL15 or LIF were significantly elevated in the mock-compared to the MCMV^S^-vaccinated animals after SARS-CoV-2 Delta (Fig. 5B, left) and Omicron BA.1 (Fig. 5B, right) challenge. As expected, expression was higher among the animals challenged with the Delta variant, than the Omicron, among mock-immunized animals. While expression was similarly reduced among MCMV^S-^immunized mice. Contrarily to inflammatory markers, T cell markers associated with the activation, differentiation, proliferation, and memory formation of T-cell responses post-vaccination were up-regulated in MCMV^S^ immunized mice compared to their controls upon Delta and Omicron BA.1-infection (Fig. 5C). In summary, our findings indicate that vaccination with MCMV^S^ not only leads to a strong immune response but also protects against sub-clinical inflammation caused by the antigenically distinct Omicron variant.

## Discussion

Several vaccines have been developed to combat the COVID-19 pandemic. Nevertheless, immune waning remains an unresolved challenge (6-8, 46), and vaccines resulting in lasting and broad protection against SARS-CoV-2 variants remain unavailable. Viral vectors that were authorized early in the COVID-19 pandemic, such as Vaxzevria (ChAdOx1-S) and the COVID-19 vaccine from Janssen (Ad26.COV2.S) (47, 48), provide immune protection that is relatively short-lived, which also afflicts the mRNA-based vaccines (49). CMV has received attention as a potential vaccine vector tool because it induces strong and lasting T-cell immunity (26). While HCMV seroprevalence is estimated to exceed 90% in some geographic areas, the pre-existing immunity to CMV does not hinder vaccine responses and protection in studies with rhesus monkeys (27, 50), arguing that CMV vectors may be used in CMV seropositive populations. MCMV shares structural and functional homologies with HCMV and allows *in vivo* analyses in the natural host. We show here, using the murine CMV vaccine vector model, that CMV-based vectors provide durable and protective immunity upon a single immunization. Hence, our data may indicate that a CMV based vaccine may provide durable and broad immune protection against COVID-19.

Neutralizing antibody responses correlate with vaccine effectiveness (51) and MCMV^S^ elicits strong neutralizing antibody responses whose affinity and neutralizing capacity increase over time (17). Here, we show that MCMV^S^ induces antibodies that neutralize the antigenically distinct Beta and Omicron variants and protect against disease upon infection with these viruses, with neutralization titers increasing for at least 6 months post-vaccination (Fig. 3B; 2J). This feature is unique for this vector and superior to other vaccine formulations, which require repeated booster administrations to avoid immune waning over time (47, 48, 52, 53). Considering that neutralizing antibodies against SARS-CoV-2 wane at similar rates upon infection or vaccination with currently used vaccines (9, 46, 54-56), our vaccine might provide an immunity that is even better than that induced by infection.

Interestingly, even the relatively weak neutralizing antibody responses against the SARS-CoV-2 Omicron variant were associated with protection against illness and lowered viral loads in the lungs (Fig. 2). One may speculate that T cell immunity elicited by MCMV^S^ vaccination protected these animals. Similar phenomena were documented in preclinical studies with rhesus macaques or in clinical studies, where study subjects were protected although no neutralizing antibodies were observed (57-59). However, we cannot exclude the possibility that memory B lymphocytes in MCMV^S^-immunized mice responded to the challenge by rapidly generating neutralizing antibodies upon challenge, thus limiting virus replication and protecting the host. Finally, it may also be possible that the concerted action of these two lymphocyte lineages protected the vaccinated mice against challenge. To differentiate between these scenarios, one may use MCMV vectors eliciting only T cell responses against S epitopes, or mice lacking T cells. However, such experiments would go beyond the scope of this study.

In this study, we showed that even an S antigen that cannot bind to the target ACE2 receptor in vaccinated subjects may still provide robust immune protection. Namely, the S antigen from the prototypical Wuhan strain of SARS-CoV-2, which was cloned into our vaccine vector, cannot bind to the murine ACE2 receptors (60). Nevertheless, MCMV^S^ protected BALB/c mice against variants that encode an N501Y mutation and thus engage the mACE2 and may replicate in mice. Some studies reported that the binding of the S1 subunit of the S protein to ACE2 on endothelial cells may affect endothelial barrier integrity and cardiac activity (61-63), which would be excluded in such approaches. While we observed no adverse effects in hACE2 mice immunized in the long term with our vector, arguing against such adverse effects (or other side effects, unrelated to ACE2 biding), our data demonstrate that spike formulations may protect against SARS-CoV-2 infection in absence of ACE2 receptor engagement by the vaccine-encoded spike, which was a concern in light of low levels of S antigen persistence in the host.

With this study we clearly show that even in Omicron infected mice, which do not display overt disease pathogenesis, sub-clinical inflammation is reduced by an MCMV vector encoding an antigenically distinct spike variant (Fig 4-5). However, Bulk RNA sequencing in inflammation research involves analyzing a mixed population of cells together, providing only a snapshot of the average genetic expression. Hence, our analysis may help to identify gene expression patterns associated with inflammation across a tissue sample, but it lacks the ability to pinpoint specific cellular contributions to gene expression profiles. Single-cell sequencing offers a more detailed view of individual cell responses in inflammation studies, allowing for a deeper understanding of cellular heterogeneity and potential therapeutic targets. Such analysis may be performed in the future to better understand the mechanisms by which immunization prevents hyper-inflammatory responses to infection.

The data presented here demonstrate that an MCMV-based vaccine candidate expressing the full-length prototype S protein is highly immunogenic and protective in mice. We demonstrated that a single dose of the MCMV^S^ may protect not only young, but also aged mice against a lethal dose of the SARS-CoV-2, and that the protection acts against immune-evasive Beta and Omicron variants. Our approach provided evidence for lifelong immunity, especially in terms of T cell responses. Future research needs to focus on a head-to-head comparison of MCMV-based vaccines with other COVID-19 vaccine formulations, on the length of protection and administration routes, paving the way towards clinical trials with CMV vectors. We propose a radical departure from the conventional vaccines against COVID-19 that provide only short-term protection that require continuous boosters and frequent adaptations of the antigen in the vaccine to circulating spike variants. This study demonstrates that in principle, such approach is feasible.

## Material and Methods

### Cell culture and viruses

Vero E6 (CRL-1586) and M2-10B4 cells (ATCC CRL-1972) were cultured as described previously (17). Caco-2 cells (ACC 169) were purchased from DSMZ (Braunschweig, Germany) and were cultured in DMEM (Gibco, NY, USA) supplemented with 20% fetal bovine serum (FBS), 2 mM L-glutamine, 100 IU/mL penicillin and 100 μg/mL streptomycin. The FI strain of SARS-CoV-2 (GISAID database ID: EPI_ISL_463008) was described previously as a D614G variant (64) and was passaged on Caco-2 cells in the biosafety level 3 (BSL3) laboratory at HZI. A clinical isolate of the SARS-CoV-2 B.1.617.2 (Delta) variant was isolated at the Fran Mihaljevic clinical center in Zagreb and propagated as previously described (64).

SARS-CoV-2 genome sequences are available on GISAID and GenBank under the following accession numbers: SARS-CoV-2 B.1.351 (Beta) FFM-ZAF1/2021 (GenBank ID: MW822592) (65) and SARS-CoV-2 B.1.1.529 (BA.1) FFM-ZAF0396/2021 (EPI_ISL_6959868; GenBank ID: OL800703) (66). MCMV^WT^ refers to the BAC-derived molecular clone of MCMV (pSM3fr-MCK-2fl clone 3.3) (67). MCMV^S^ was generated by *en passant* mutagenesis, as described previously (17). Briefly, the codon-optimized S protein of the Wuhan variant replaced the coding sequence of the viral ie2 protein.

The expression vector for SARS-CoV-2 S protein of Omicron (BA.1) (based on isolate hCoV-19/Botswana/R40B58_BHP_3321001245/2021; GISAID Accession ID: EPI_ISL_6640919), BA.4/5 (based on isolate hCoV-19/England/LSPA-3C01A75/2022 was generated by Gibson assembly as described previously (68) and then subsequently introduced in pseudotype VSV backbone that lacks VSV glycoprotein G (VSV-G) (69). A plasmid encoding the S protein of SARS-CoV-2 Beta (B.1.351) has been previously reported (17).

### Virus stock generation and plaque assay

BAC-derived MCMV was reconstituted by transfection of BAC DNA into NIH-3T3 cells (ATCC CRL-1658) using FuGENE HD transfection reagent (Promega, WI, USA) according to the manufacturer’s instructions. Transfected cells were cultured until viral plaques appeared and passaged 5 times in M2-10B4 cells before virus stock production. Virus stocks were prepared on ice. First, supernatants of infected M2-10B4 cells were collected and infected cells were pelleted (5,000 × g for 15 min). The resulting cell pellets were homogenized in DMEM supplemented with 5% FBS and cell debris was removed by centrifugation (12,000 x g for 10 min). Collected supernatants were resuspended in VSB buffer (0.05 M Tris-HCl, 0.012 M KCl, and 0.005 M EDTA, adjusted to pH 7.8) and then concentrated by centrifugation through a 15% sucrose cushion in VSB buffer (23,000 × g for 1.5 h). The resulting pellet was resuspended in 1-1.5 mL VSB buffer, briefly spun down, and supernatants were aliquoted and kept at -80°C. BAC-derived mutant MCMV^S^ were propagated on M2-10B4 cells and concentrated by sucrose density gradient centrifugation.

SARS-CoV-2 D614G was generated and viruses were quantified by plaque assays as described before (17) with a minor modification that Caco-2 cells were used for virus production. SARS-CoV-2 Beta, Delta and Omicron BA.1 stocks were generated as described previously and titers were determined by the median tissue culture infective dose (TCID50) method (66).

Pseudotyped viruses were harvested as described before (17, 68). In brief, 293T cells were transfected with expression plasmids (pCG1) encoding different S proteins of SARS-CoV-2 variants by using the calcium-phosphate method. At 24h post-transfection, the medium was removed and cells were inoculated with a replication-deficient VSV vector lacking its glycoprotein and coding instead for an enhanced green fluorescent protein (GFP) (kindly provided by Gert Zimmer, Institute of Virology and Immunology, Mittelhäusern, Switzerland). Following 1 h incubation at 37°C, the cells were washed with PBS, and culture media containing anti-VSV-G antibody (culture supernatant from I1-hybridoma cells; ATCC CRL-2700) were added. The pseudotype virus was harvested at 16-18 h post-infection.

### Virus in vivo infection

K18-hACE2 mice were obtained from Jackson Laboratories and bred in the core animal facility of Helmholtz Center for Infection Research, Braunschweig. BALB/c mice were purchased from Envigo (IN, USA). All animals were housed under Specific Pathogen Free (SPF) conditions at HZI during breeding and infection. All animal experiments were approved by the Lower Saxony State Office of Consumer Protection and Food Safety, license number: 33.19-42502-04-20/3580.

K18-hACE2 mice (2-6 months old and >17 months) were intraperitoneally (i.p.) immunized with 10^6^ PFU of recombinant MCMV^S^ or MCMV^WT^ diluted in PBS or treated with PBS (200 μl i.p. per animal). BALB/c mice (4-7 months old) were i.p. immunized with 2×10^5^ PFU of recombinant MCMV^S^ or MCMV^WT^. All mice were weekly or bi-weekly monitored and scored for their health status after vaccination. Blood was isolated at indicated time points by puncture of the retrobulbar plexus or by cardiac bleeding upon death. SARS-CoV-2 challenge experiments were performed in the HZI BSL3 laboratory essentially as described (70) with the following modifications: K18-hACE2 mice were intranasally (i.n.) infected with 2×10^3^ PFU of D614G, Delta or Omicron BA.1 SARS-CoV-2 variant, while BALB/c mice were challenged with 6×10^4^ PFU of the VoCs. SARS-CoV-2-infected mice were monitored for body mass loss and clinical status at least daily, according to the animal permit.

SARS-CoV-2-challenged animals were scored daily to monitor any signs of disease development. Animals were scored based on five criteria: spontaneous/social behaviour, fur, fleeing behaviour, posture, and body mass loss. Each score indicates the following: no signs of symptoms (score=0), mild and/or sporadic symptoms (score=1), moderate and/or frequent symptoms (score=2), and severe symptoms with a clear sign of heavy suffering (score=3). Weight loss criterion is scored as follows: ≤1 % (score=0), 1-10 % (score=1), 10-20% (score=2), and >20% (score=3). Mice with a score of 3 in one criterion, or an overall score of ≥9, were removed from the experiments.

### Organ harvest

Animals were euthanized by CO_2_ inhalation, blood was collected from the heart and kept at room temperature (RT) for 30–60 min until clotting occurred, then stored at 4 °C. For further analysis, serum was isolated by centrifugation. The trachea, lungs, heart, spleen, stomach, and brain were harvested at 3, 5 or 7 days post-SARS-CoV-2 challenge. In animals that reached humane endpoints before 7 days post-challenge, organs were harvested on the day of euthanasia. Solid organs were weighed and homogenized in bead-containing lysis tubes (Lysis tube E, Analytik Jena) in 500 μL or 1,000 μL PBS with an MP Biomedical FastPrep 24 Tissue Homogenizer (MP Biomedicals, CA, USA) (full speed, 2 × 20 s) and stored at -80°C. Samples designated for RNA-isolation or bulk sequencing were mixed with Trizol reagent (Invitrogen) at a 1:3 ratio. All centrifugation steps were carried out at 10,000× g for 10 min at 4 °C.

### RNA isolation and viral loads analyses

RNA isolation was performed with the Rneasy RNA isolation kit (Quiagen) or innuPrep Virus TS RNA kit (Analytik Jena), according to the manufacturer’s protocol. Shortly, for the first experiments of infected K18-hACE2 mice challenged with D614G the Rneasy RNA isolation kit was used by adding 250 μL of organ homogenates in 750 μL Trizol and subsequently centrifuged at 16,000 x g for 3 min. The resulting supernatants were carefully collected and washed with the same volume of 70% Ethanol. The mixed solution was transferred into a collection tube and centrifuged at 10,000 x g for 30 sec. After decanting the flow-through, the column was washed once with 700 μL of RW1 wash buffer and twice with 500 μL RPE buffer. Lastly, 40 μL of nuclease-free water was added to the column for RNA elution. For the remaining experiments including the BALB/c and K18-hACE2 mice challenged with the SARS-CoV-2 VoCs RNA extraction was conducted using the innuPrep Virus TS RNA kit (Analytik Jena) following the manufacturer’s instructions. In brief, Trizol-inactivated samples were combined with an equal volume of lysis solution CBV, which contained carrier RNA and Proteinase K. This mixture was then incubated at 70 °C for 10 minutes. After the lysis step, samples were blended with two sample volumes of isopropanol. Subsequently, the samples were applied to the provided spin filters, washed with washing buffer (LS), and rinsed with 80% ethanol. RNA was finally eluted in 60 μL of RNase-free water and stored at -80 °C after determining the RNA concentration using a NanoDrop (Thermo Scientific NanoDropOne).

Eluted RNAs were analyzed further to assess viral RNAs in the given organs by quantitative reverse transcription polymerase chain reaction (RT-qPCR). The reaction was performed with a total volume of 20 μL containing 2 μL of sample RNAs or positive control RNAs, 5 μL TaqPath 1-step RT-qPCR Master Mix with ROX reference dye, and 1.5 μL probe/primer sets. 2019-nCoV RUO kit was used to detect SARS-CoV-2 RNAs (Integrated DNA Technologies (IDT), USA), and Taqman Rodent GAPDH control reagents (ThermoFischer Scientific, USA) were used for endogenous GAPDH RNAs. For absolute viral RNA quantification, a standard curve was generated by serially diluting a SARS-CoV2 plasmid with the known copy numbers 200,000 copies/μL (2019-nCoV_N_Positive Control, #10006625, IDT, USA) at 1:2 ratio in all PCR analyses, with a quantitation limit of 20 copies of the plasmid standard in a single qPCR reaction. The viral RNA of each sample was quantified in triplicate and the mean viral RNA was calculated by the standard. RT-qPCR was performed using the StepOnePlusTM Real-Time PCR system (ThermoFischer Scientific, USA) according to the manufacturer’s instructions. SARS-CoV-2 N-gene copy numbers were normalized to total RNA input and to rodent GAPDH copy numbers.

### Detection of infectious SARS-CoV-2 titers by plaque assay

Lung and brain organ homogenates were serially diluted in DMEM (Gibco, NY, USA) supplemented with 5% FBS and 100 IU/mL penicillin, and 100 μg/mL streptomycin. 100 μL of sample dilutions were transferred onto confluent VeroE6 cells in a 96-well format. After inoculation for 1 h at 37 °C, the inoculum was removed and 1.5% methylcellulose in MEM supplemented with 5% FBS, 2 mM L-glutamine, 100 IU/mL penicillin, and 100 μg/mL streptomycin was added to the cells. The infected cells were incubated at 37 °C for 48 h before inactivation with a 4% formalin solution in PBS for 10 min at RT. The fixed cells were subjected to immunofluorescent staining against the SARS-CoV-2 N protein. Briefly, fixed cells were permeabilized with 0.1% Triton X-100 (Sigma-Aldrich, MA, USA) for 10 min at RT and blocked with 1% BSA (Sigma-Aldrich, MA, USA) in PBS for 30 min at RT. Thereupon, cells were incubated with a monoclonal anti-SARS-CoV-2 N protein antibody (Abcalis, AB84-E02, 10 μg/mL) for 30 min at RT. After washing three times with PBS with 0.05% Tween-20 (PBS-T), a secondary antibody anti-mouse IgG conjugated with Alexa488 (Cell Signaling Technology, #4408, 1:500) was added for 30 min at RT. After washing three times with PBS-T, the stained cells were visualized using Incucyte S3 (Sartorius; GUI software versions 2019B Rev1 and 2021B).

### Flow cytometry quantification of S-specific T cells

Peripheral blood was harvested and red blood cells were removed by short osmotic shock. Thereupon, lymphocytes were stained with S-derived VNFNFNGL-specific tetramers (Kindly provided by Ramon Arens, Leiden University) for 30 min at RT. Subsequently, cells were stained with fluorescent-labeled antibodies against CD3 (17A2, eBiosciences, CA, USA), CD4 (GK1.5, BioLegend, CA, USA), CD8a (53-6.7, BD Bioscience, 528 CA, USA), CD44 (IM7, BioLegend, CA, USA), CD11a (M17/4, BioLegend, CA, USA), CD62L (MEL-14, BioLegend, CA, USA), CD127 (SB/199, Biolegend, CA, USA) and KLRG1 (2F1, BioLegend, CA, USA) for 30 min at 4°C. Dead cells identified by 7-AAD viability staining solution (BioLegend, CA, USA) were excluded from all analyses. The labeled cells were analyzed by flow cytometry (BD LSRFortessaTM Cell Analyzer) and subsequent analyses were done in detail in FlowJo Software v10.

### Detection of anti-spike antibodies in mouse sera

ELISA (Enzyme-linked ImmunoSorbent Assay) was used to detect SARS-CoV-2 Spike specific IgG response in mouse sera after 6, 12, 24 and 64 weeks post-immunization. As control, sera from mock-immunized mice were included. S1-S2 wildtype and S1-S2 B.1.1.529 variant (Omicron) were used as antigens, which were produced in insect-cells as described before (6, 70), in a baculovirus-free system. For antigen coating, 100 ng/well S1-S2 in carbonate buffer (50 mM NAHCO_3_/Na_2_CO_3_, pH 9.6) was immobilized overnight at 4 °C in 96x well-plates (Greiner Bio-One). ELISA plates were blocked with 2% MPBS-T (2% (w/v) skim milk powder, 0.05% Tween20 in PBS) for 1 h at RT and washed with H_2_O containing 0.05% Tween20 using EL405 S washer (BioTek). Mice sera were diluted 1:100 and titrated to a final dilution of 1:6400 and incubated for 1h at RT. After washing as described above, the serum IgG respone was detected using horseradish peroxidase (HRP) conjugated goat-anti mouse serum (A0186, Sigma) (1:42000 diluted in 2% MPBS-T) and plates washed again after incubation at RT for 1h. Binding was visualized using tetramethylbenzidine (TMB) substrate (19 parts TMB A solution (30 mM Potassium citrate; 1% (w/v) citric acid (pH 4.1)) and 1 part TMB B solution (10 mM TMB; 10% (v/v) Acetone; 90% (v/v) Ethanol; 80 mM H_2_O_2_ (30%))) and developed for 15 min at RT. Reaction was stopped using 0.5 M/1N sulfuric acid and absorbance measured at 450 nm with subtracting reference at 620 nm using ELISA plate reader (Epoc, BioTek). For K18-hACE2 mice challenged with D614G for unspecific binding. EC50 was analyzed by a statistical analysis tool in GraphPad Prism 9. For the aged K18-hACE2 mice challenged with the SARS-CoV-2 VoCs the Area under the curve (AUC) was calculated.

### Antibody avidity assay

Antibody avidity assay was performed as described by *Welten et. al*. 2016 (27). In brief, 100 ng/well S1-S2 were immobilized in carbonate buffer overnight at 4° and plates blocked with 2% MPBS-T for 1h at RT. Plates were washed with H_2_O containing 0.05% Tween20 using EL405 S washer (BioTek). Mice sera showing the best IgG response were selected and always three sera pooled and diluted 1:300 in 2% MPBS-T. After incubation at RT for 1h, plates were washed as described above followed by incubation with NaSCN at RT in indicated dilutions. As control, binding without NaSCN was included. After 15 min incubation, plates were washed immediately two times to ensure complete removal of NaSCN. Binding was detected using goat-anti mouse serum (A0168, Sigma) conjugated with HRP (1:42000 dilution in 2% MPBS-T) at RT for 1h and plates washed as described above. Antibody binding was visualized using TMB substrate as described above and reaction stopped after 15 min at RT using 0.5M/1N sulfuric acid. Absorbance was measured as described above and IgG binding normalized to 100% in absence of NaSCN and avidity calculated as binding after treatment with indicated NaSCN dilutions on described antigens.

### In vitro live virus neutralization titer (VNT) assay

The serum neutralization assay was performed as described before (17). Briefly, heat-inactivated sera were serially diluted and incubated with 100 PFU/100 μL of SARS-CoV-2 for 1 h at RT. Thereupon, they were transferred to 96-well plates seeded with Vero-E6 cells and inoculated with serum and virus for 1 h. After the inoculum removal, the cells were overlaid with 1.5% methylcellulose and incubated at 37°C and 5% CO_2_ for 3 days. The cells were fixed with 4% formaldehyde, followed by crystal violet staining and plaque counting. Serum-neutralizing titer that results in a 50% reduction of virus plaques (VNT50) was analyzed by GraphPad Prism 9 nonlinear regression analysis.

### Pseudovirus neutralization assay

Pseudovirus neutralization assays were performed as described in the previous publications (17, 71). For neutralization experiments, pseudotyped particles and heat-inactivated serum dilution were mixed at a 1:1 ratio and incubated for 60 min at 37°C before being inoculated onto VeroE6 cells grown in 96-well plates. At 24 h post-infection, GFP expression was measured by using Incucyte S3 (Sartorius, Goettingen, Germany).

### bulk RNA extraction and sequencing

To perform RNA bulk sequencing, RNA was isolated from lung tissue and blood using Trizol reagent (Invitrogen) at a 1:3 ratio. Briefly, 3x Trizol LS was added to the homogenized organ sample or blood and vortexed thoroughly. After incubation, 1/5 of the volume of chloroform were added. The samples were vortexed again and incubated for 5 min at RT. Subsequently, tubes were centrifuged at 12,000 × *g* for 15 min at 4°C and aqueous phase were transferred into a new tube and RNA was extracted with the RNA Clean and Concentrator kit (ZYMO Research).

Bulk RNA sequencing libraries were constructed using the NEBNext Ultra II Directional RNA Library Prep Kit (New England Biolabs) and sequenced on a high throughput NextSeq 500 device. Reads were aligned to the Mus musculus genome (GRCm39 or mm39) using hisat2 (72) and gene expression quantified using the package featureCounts from Rsubread (73). Analysis was done with DESeq2 (74). Differentially expressed genes were defined by an absolute fold change in mRNA abundance greater than 1.5 (log2 fold change of 0.58 - using DESeq2 shrunken log2 fold changes) and an adjusted p-value of less than 0.05 (Benjamini-Hochberg corrected). Gene expression deconvolution was performed with the package GEDIT, using the reference matrix provided for blood analysis (75).

### Statistics

One-way ANOVA with Kruskal-Wallis correction was performed for multiple-group analysis. Two-tailed Mann-Whitney tests were used to compare the difference between two independent groups. Log-rank (Mantel-cox) tests were performed to compare the survival distributions of groups. Statistical analysis was calculated by GraphPad Prism 9.

## Data availability

All SARS-CoV-2 genome sequences are available as mentioned above. The D614 SARS-CoV-2 variant (GISAID database ID: EPI_ISL_463008), SARS-CoV-2 B.1.351 (Beta) FFM-ZAF1/2021 (GenBank ID: MW822592), and SARS-CoV-2 B.1.1.529 (BA.1) FFM-ZAF0396/2021 (EPI_ISL_6959868; GenBank ID: OL800703).

## Author Contributions

Conceptualization, KM, HJ, YK and LCS; methodology, KM, HJ, YK, LGTA, UK, KE, MZC, MH, FB, MR, MHu, MCB, JM, MS, AK; validation, LCS, ML, SP, MHu; formal analysis, KM, HJ, YK, LGTA; investigation, KM, HJ, YK, LGTA; resources, LCS, ML, SP, MHu, MW, SJ and SC; data curation, KM, HJ, YK, LGTA; writing—original draft preparation, KM and YK; writing—review and editing, all authors; supervision, LCS; project administration, LCS; funding acquisition, LCS, SJ and ML

## Institutional Review Board Statement

All animal experiments were approved by the Lower Saxony State Office of Consumer Protection and Food Safety, license number: 33.19-42502-04-20/3580.

## Acknowledgments

We would like to thank Bettina Fuerholzner, Tatjana Lueddecke, Leila Abassi and Daniela Lenz for their excellent help with the animal experiments, as well as Ayse Barut and Inge Hollatz-Rangosch for their technical assistance. We acknowledge colleagues from HZI for their professional expertise, Lothar Gröbe from the flow cytometry facility, Susanne Talay from the S3 facility, Marina Pils, Katrin Schlarmann, Petra Beyer, and Bastian Pasche from the animal facility, and Katarzyna M. Sitnik and Natascha Goedecke for support with animal ethical issues.

