## Supplementary Material for "A single-dose MCMV-based vaccine elicits long-lasting immune protection in mice against distinct SARS-CoV-2 variants"

### Supplementary Figures and Figure legends

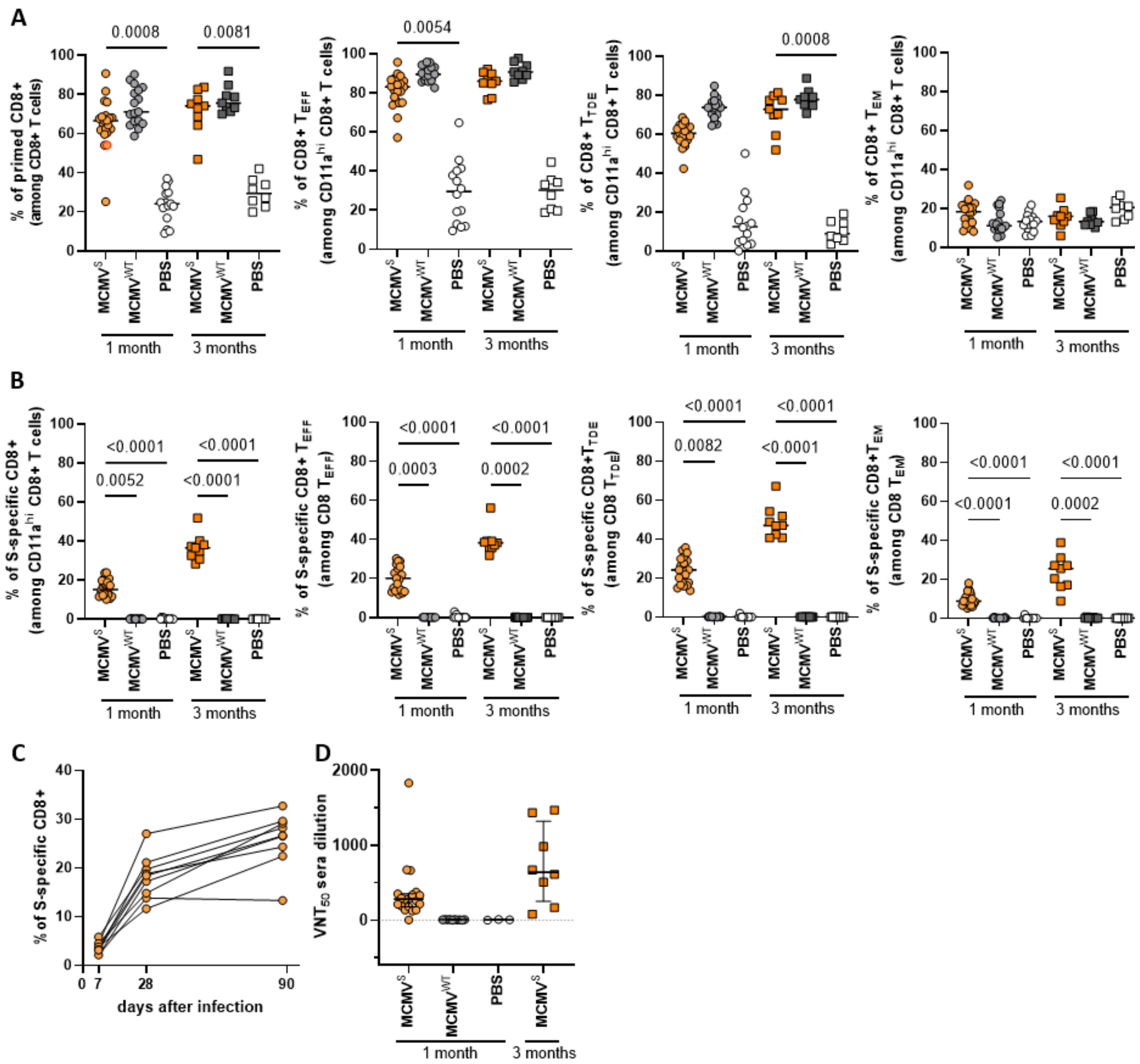


**Supplementary Figure 1: Protection against SARS-CoV-2 D614G elicited by a single dose of $\text{MCMV}^{S}$ vaccination.** (A) CD3+CD8+CD4- T cells were gated for the primed subpopulation ($CD11^{hi}CD44^{hi}$ - *leftmost panel*). Primed CD8 T cells were progressively gated into $T_{EFF}$ ($CD62L^{lo}$), $T_{TDE}$ ($CD62L^{lo}KLRG1^{hi}$), or $T_{EM}$ ($CD62L^{lo}KLRG1^{lo}$) subpopulations. Frequencies of cells in each subset as a fraction of the parental population are shown. (B) The percentages of S-specific cells in each subset shown in (A) are displayed. (A-B) Each symbol indicates an individual mouse and each color represents vaccination with following vectors: Orange=$\text{MCMV}^{S}$, Grey=$\text{MCMV}^{WT}$, and White=PBS. Horizontal lines show geometric means. Kruskal-Wallis tests were used for the statistical analyses (1 month: n=21 $\text{MCMV}^{S}$, n=15 $\text{MCMV}^{WT}$, n=17 PBS; 3 months: n=9 $\text{MCMV}^{S}$, $\text{MCMV}^{WT}$, n=8 PBS). (C) Antigen-specific memory responses over time in blood shown as the percentage of antigen-specific cells within the total CD8 compartment. Each line connects values from an individual mouse at indicated time points. (D) Violin plots of virus neutralization titers (VNTs) that resulted in a 50% of reduction of SARS-CoV-2 infection (VNT50). Each symbol indicates an individual mouse.

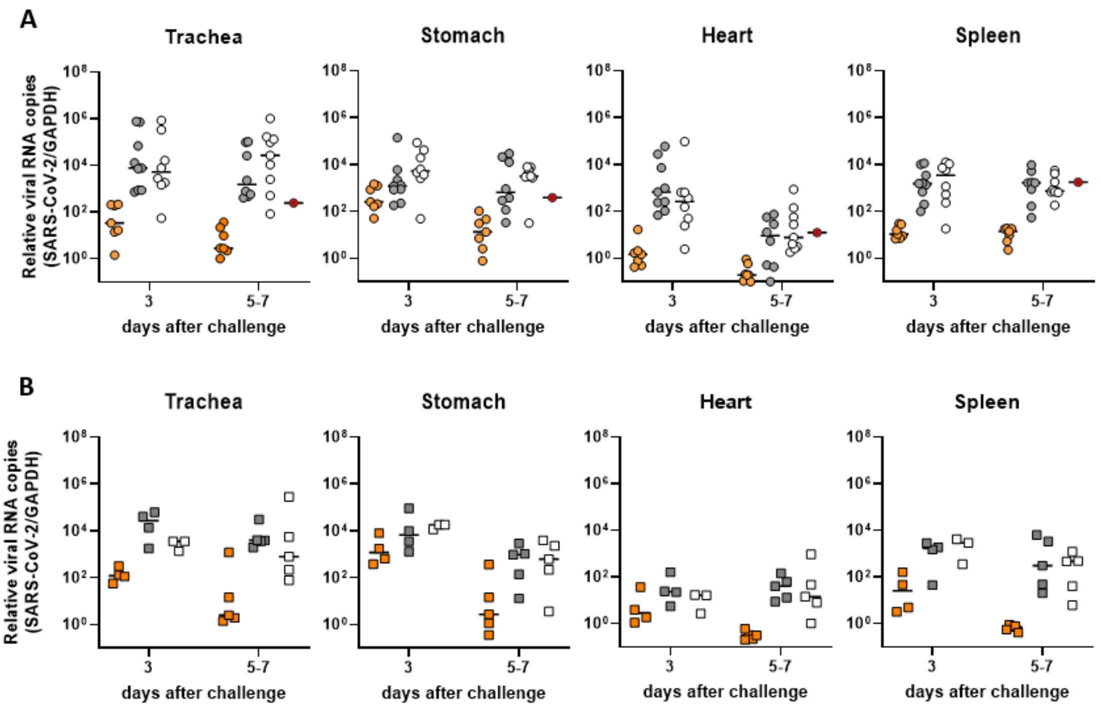


**Supplementary Figure 2: Protection against SARS-CoV-2 D614G elicited by a single dose of MCMV$^{S}$ vaccination. (A-B)** Relative viral RNA loads in the representative organs (trachea, stomach, heart, spleen) of **(A)** 6- and **(B)** 12-weeks immunized mice. Each symbol indicates an individual mouse and each color represents vaccination with following vectors: Orange=MCMV$^{S}$, Grey=MCMV$^{WT}$, and White=PBS. Organs were harvested at the indicated time points of 3 and 5-7 days after SARS-CoV-2 D614G challenge. SARS-CoV-2 viral RNA was normalized to the housekeeping gene mGAPDH. Horizontal lines indicate medians. Crossed red symbols indicate no animals alive by the time of analyses. Organs of mock-immunized mice that reached humane end-points before 7 days upon challenge were harvested on the day of euthanasia. Pooled data (n=6-8 per group) from two independent experiments are shown.

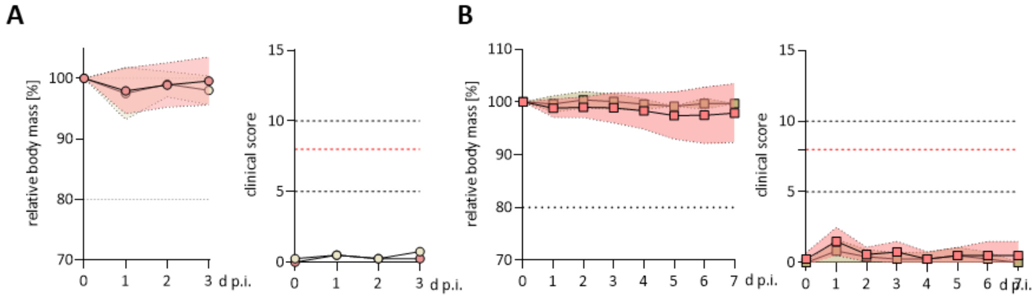


**Supplementary Figure 3: Sustained neutralizing antibody responses against Beta and the Omicron variants upon MCMV$^{S}$ immunization. (A, B)** Percentage of body weight (right) and clinical score (left) upon challenge are shown. Mice were challenged with $2x10^{3}$ PFU of SARS-CoV-2 Omicron BA.1 6 weeks **(A)** or 12 weeks **(B)** post immunization. The red dotted line indicates the threshold in body weight loss resulting in a humane end-point.

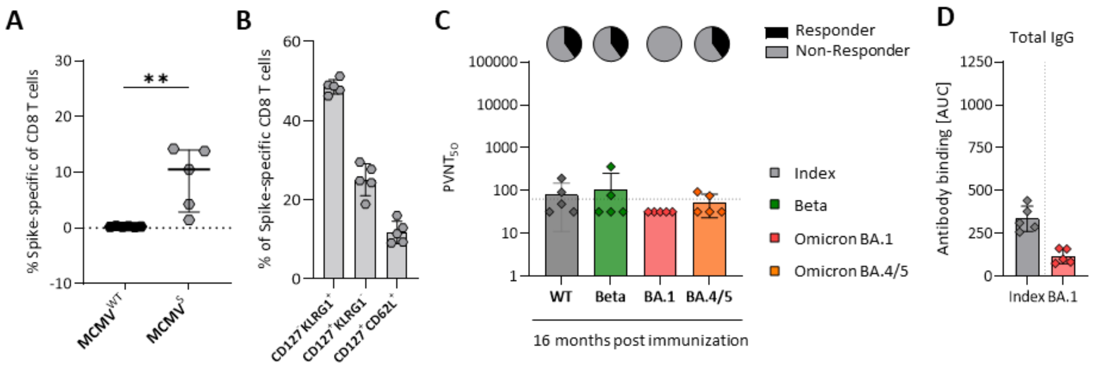


**Supplementary Figure 4: Single-dose of MCMV$^S$ provides inflationary immunogenicity in aged mice.** Immune response in aged K18-hACE2 mice 16 months after immunization with 2x10$^5$ PFU/ml MCMV$^S$. **(A)** Dot plot displays median of percentage of the frequency of Spike-specific CD8 T cells in the spleen. Each data point represents a single mouse (MCMV$^{WT}$: n= 6; MCMVS: n=5). **(B)** Primed CD8 T cells were progressively gated into Effector-memory cells ($T_{EM}$): CD8$^+$/CD44$^+$/Tet$^+$/CD127$^+$/KLRG1$^-$, Effector-like cells: CD8$^+$/CD44$^+$/Tet$^+$/CD127$^-$/KLRG1$^+$ or Central memory cells ($T_{CM}$): CD8$^+$/CD44$^+$/Tet$^+$/CD127$^+$/CD62L$^+$. (C) Pseudovirus neutralization titers (PVNTs) that resulted in a 50% of reduction of SARS-CoV-2 infection (PVNT50). Sera was tested against the Index (Wuhan; *grey*), Beta *(blue)*, Omicron BA.1 *(red)* and Omicron BA 4/5 *(orange)* SARS-CoV-2 strain. Each symbol indicates an individual mouse. Solid horizontal lines show the median and the dotted line indicates the detection limit. Pie charts indicate the percentage [%] of animals, that responded *(black)* or non-responded (grey) with neutralizing antibody titers. **(D)** Amount of SARS-CoV-2 Spike specific total IgG response against the Index (Wuhan) and Omicron BA.1 VoC in mouse sera

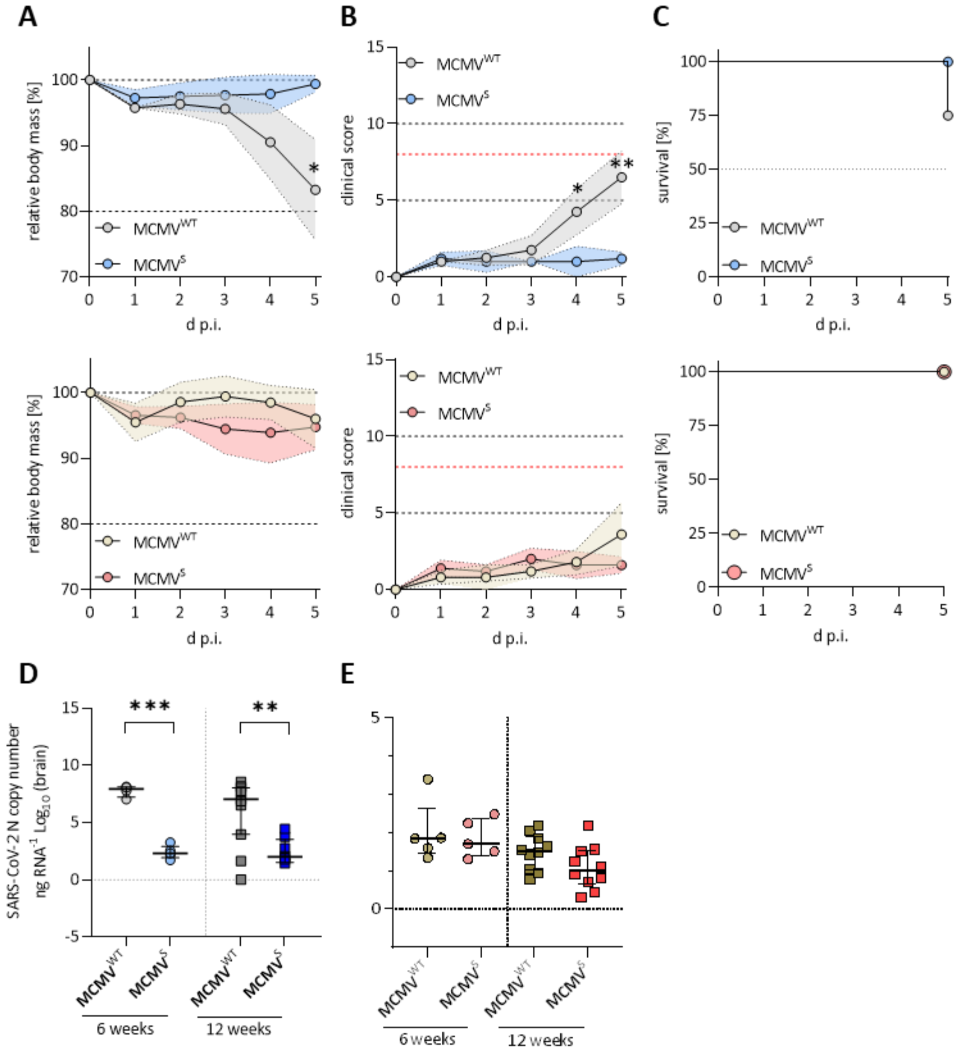


**Supplementary Figure 5: Aged MCMV$^{S}$ immunized mice show full and long-lasting protection against heterologous SARS-CoV-2 challenge. (A-C)** Mice were challenged with $2x10^3$ PFU of SARS-CoV-2 Delta or Omicron BA.1 after 6 weeks immunization. **(A)** Percentage of body weight and **(B)** daily clinical scores upon challenge with Delta (*top*) or Omicron BA.1 (*bottom*) are shown. The red dotted line indicates the threshold in score resulting in a humane end-point. **(C)** Survival kinetics of challenged mice representing the Percentage [%] of mice reaching a humane end point. A log-rank (Mantel-cox) test was used for the statistical analysis. **(D-E)** SARS-CoV-2 N gene copy numbers as copy number/ng RNA Log10 at 5 dpi with SARS-CoV-2 **(D)** Delta or **(E)** Omicron BA.1 in brain homogenates Statistical significance for was calculated using Welch's t test (two-tailed) ( ** p <0.01, *** p < 0.001).
